# Identification of a core module for bone mineral density through the integration of a co-expression network and GWAS data

**DOI:** 10.1101/803197

**Authors:** Olivia L Sabik, Gina M Calabrese, Eric Taleghani, Cheryl L Ackert-Bicknell, Charles R Farber

## Abstract

Recently, the “omnigenic” model of the genetic architecture of complex traits proposed two general categories of causal genes, core and peripheral. Core genes are hypothesized to play a direct role in regulating disease; thus, their identification has the potential to reveal critical regulators and novel therapeutic targets. Here, we sought to identify genes with “core-like” characteristics for bone mineral density (BMD), one of the most significant predictors of osteoporotic fracture. This was accomplished by analyzing genome-wide association study (GWAS) data through the lens of a cell-type and timepoint-specific gene co-expression network for mineralizing osteoblasts. We identified a single co-expression network module that was enriched for genes implicated by GWAS and partitioned BMD heritability, correlated with *in vitro* osteoblast mineralization, and enriched for genes, which when mutated in humans or mice, led to a skeletal phenotype. Further characterization of this module identified four novel genes (*B4GALNT3*, *CADM1*, *DOCK9*, and *GPR133*) located within BMD GWAS loci with colocalizing expression quantitative trait loci (eQTL) and altered BMD in mouse knockouts, suggesting they are causal genetic drivers of BMD in humans. Our network-based approach identified a “core” module for BMD and provides a resource for expanding our understanding of the genetics of bone mass.

## Introduction

Osteoporosis is a disease characterized by low bone mineral density (BMD) and an increased risk of fracture ^1^. Worldwide, it is one of the most common diseases, affecting over 200 million individuals and causing 8.9 million fractures annually ^2^. Although osteoporosis is a multifactorial disease influenced by both environmental and genetic variation, fracture-related traits, such as BMD, are influenced, in large part, by genetics (h_2_>0.5) ^3–5^. Over the last decade large-scale genome wide association studies (GWASs) have begun to dissect the genetics basis of bone traits with a primary focus on BMD ^6, 7^. These studies have been tremendously successful, identifying over 1100 independent BMD associations ^8–10^. However, despite the wealth of genetic signals, the genes and mechanisms through which these associations impact bone remain largely unknown ^6, 7^.

Recently, the “omnigenic model” was proposed as a framework for understanding the genetic architecture of complex traits, such as BMD ^11, 12^. The model posits that all genes expressed in disease-relevant cell-types have the potential to contribute to disease variation. One of the key concepts of the omnigenic model is the classification of causal genes as either “core” or “peripheral”. Core genes are predicted to directly modulate traits; whereas, peripheral genes are expected to impact traits via their effects on networks of core genes ^12^. The distinction between core and peripheral genes is logical given the evidence demonstrating that the contributions of genes to a disease or phenotype are not equal. As an example, *RUNX2* is a transcription factor and master regulator of osteoblast activity and bone formation that initiates a transcriptional program absolutely required for the formation of a mineralized skeleton ^13^. In contrast, hundreds of genes have been identified participating in myriad pathways whose absence has subtle, often context-dependent (such as age and sex), effects on bone ^8, 9, 14^. Furthermore, we know the same distinction lies in biological processes, some of which play an intimate role in the regulation of a trait, while others play minor accessory roles. Thus, the identification of causal genes from GWAS data and the labeling of such genes as core or peripheral has the potential to highlight previously undiscovered key regulatory genes for specific trait-related biological processes, which may be more ideal therapeutic targets.

There are two main challenges in the identification of core genes. The first is how to precisely define them ^12, 15–17^. In the omnigenic model, a gene is defined as a “core” gene “if and only if the gene product (protein, or RNA for a noncoding gene) has a direct effect—not mediated through regulation of another gene—on cellular and organismal processes leading to a change in the expected value of a particular phenotype” ^11, 12^. This statistical definition is convenient for explaining the omnigenic model, but is difficult to utilize for the identification of core genes in practice. It is also very strict; e.g. is *RUNX2,* as described above, a core gene for BMD? Instead we propose to use a set of biologically motivated criteria to distinguish genes with core-like properties from those that are likely peripheral by leveraging known pathways and processes that are essential to a disease-associated trait. For example, we would expect the expression of genes with core-like properties operating in pathways of critical importance in the regulation of BMD to be correlated with BMD and their severe perturbation to have a substantial impact on BMD (e.g., monogenic disease genes).

The second challenge is designing a strategy to identify genes with core-like properties, since GWAS alone is incapable of determining whether a locus is driven by a core or peripheral gene. One of the primary tenets of the omnigenic model is that peripheral genes account for a substantial component of the heritability of a trait because their effects are amplified by interactions with networks of co-expressed core genes ^12^. If one expects core genes to be co-expressed then integrating the results of GWAS with co-expression networks, which reflect the transcriptional programs associated with the trait of interest, is a logical approach to identify modules of genes with core-like properties. A number of studies have already successfully used co-expression networks to inform GWAS, however this approach has not been used in the context of the omnigenic model ^18–22^.

Here, we combine weighted gene co-expression network analysis (WGCNA) and BMD GWAS data to identify genes that are causal genetic drives of BMD with core-like properties. Our approach used a co-expression network for mature, mineralizing osteoblasts which we hypothesized would allow us to identify core genes specific for the process of mineralization. We first identified network modules enriched for genes implicated by GWAS and partitioned BMD heritability and then used the following biologically motivated filters to identify modules enriched for genes with core-like properties (i.e. “core” modules): (1) correlation with *in vitro* mineralization (a process of fundamental importance to BMD), (2) enrichment for genes that, when knocked-out in mice, alter BMD, and (3) enrichment for monogenic skeletal disease genes. Our analysis identified a single module (referred to as the “purple” module) fulfilling all the proposed criteria of a core module. As would be expected of a core module for mineralization, the purple module was enriched for genes with well-known roles in osteoblast activity and bone formation. Furthermore, we identified two submodules of genes within the purple module that followed distinct patterns of expression across osteoblast differentiation, the early and the late differentiation submodule (EDS and LDS). We found that the LDS, relative to the EDS, was more enriched for genes with core-like properties. Supporting the hypothesis that many LDS genes are causal genetic drivers, we observed that lead BMD SNPs located in GWAS loci harboring an LDS gene were more likely to overlap active regulatory elements in osteoblasts. Further characterization of the LDS identified four novel genes (*B4GALNT3*, *CADM1*, *DOCK9*, and *GPR133)* located within BMD GWAS loci that had colocalizing human eQTL and altered BMD in mouse knockout studies. We anticipate that this integrative approach will aid in the search for genes with core-like properties and pathways underlying BMD and risk of fracture.

## Results

### I. Construction of a co-expression network reflecting transcriptional programs in mineralizing osteoblasts

The goal of this work was to use a cell- and stage-specific co-expression network to identify genes with core-like properties that are causal for BMD GWAS associations. We chose to focus on generating a co-expression network using transcriptomic data from a single cell-type at a single-time point during differentiation: mature, mineralizing osteoblasts. We hypothesized this would allow us to focus on genes with core-like properties in the context of mineralization, a process critical in the regulation of BMD. We began by using WGCNA to construct a co-expression network using transcriptomic profiles generated from mineralizing primary calvarial osteoblasts from 42 strains of Collaborative Cross (CC) mice ^23^. The CC is a panel of genetically diverse recombinant inbred strains. The resulting network consisted of 65 modules of genes, with an average of 292 genes per module (**Figure 1 and Supplemental File 1**). Each co-expression module was distinguished by its assigned color (e.g., the purple module).

**Figure 1.**
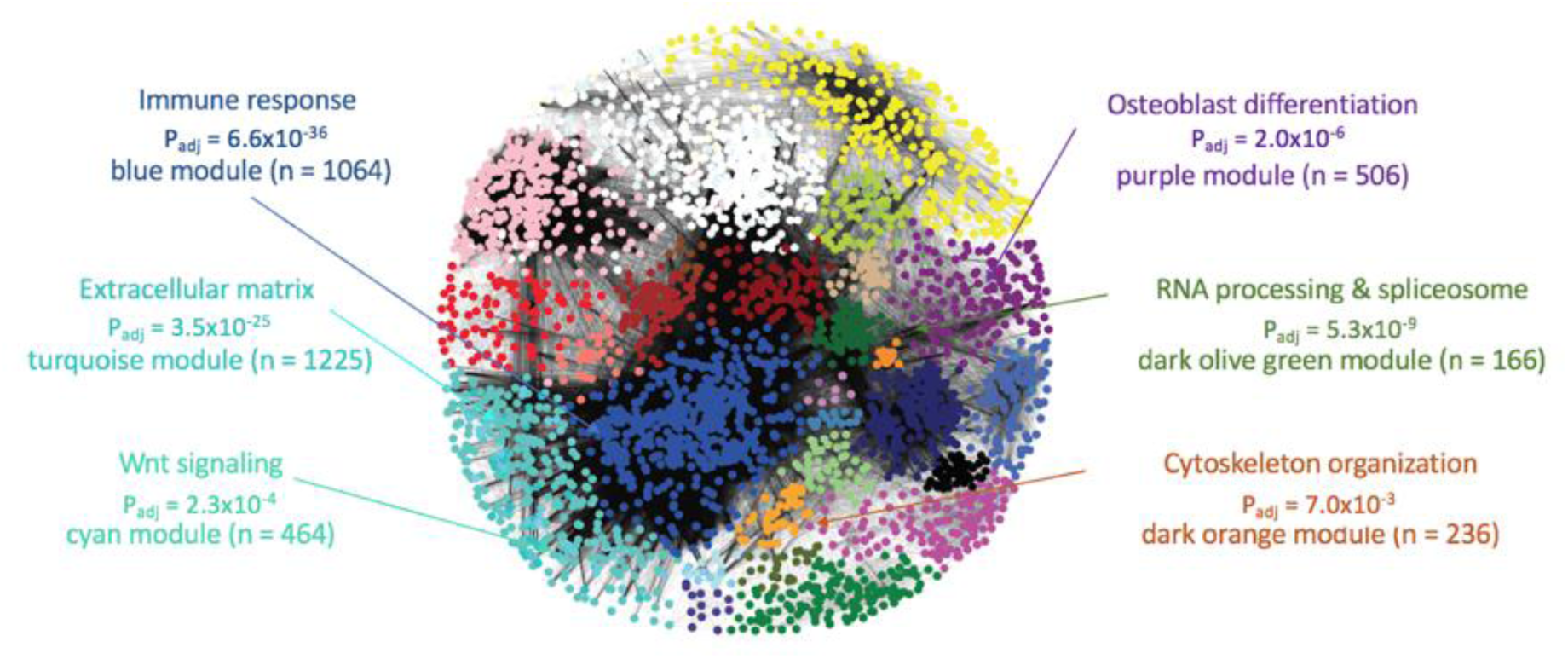
Weighted gene co-expression network generated using transcriptomic profiles from mineralizing osteoblasts. The network was composed of 65 modules of co-expressed genes, many of which were enriched for specific biological processes relevant to osteoblasts.

To confirm that modules of genes produced by the co-expression analysis represented transcriptional programs reflecting specific biological processes, we assessed whether modules were enriched for genes associated with specific gene ontology (GO) terms ^24^. Most network modules were enriched for general biological processes, such as the immune response (P_adj_ = 6.6 × 10_−36_) in the blue module, mRNA metabolism (P_adj_ = 7.8 × 10_−9_) in the darkolivegreen module, and chromatin remodeling (P_adj_ = 1.9 × 10_−4_) in the grey60 module (**Figure 1 and Supplemental File 2**). However, as would be expected, there were a subset of modules enriched for genes involved in the activity of osteoblasts. For example, the cyan module was enriched for members of the Wnt signaling pathway (a key regulator of osteoblast activity) (P_adj_ = 2.3 × 10_−4_), the turquoise module was enriched for genes encoding extracellular matrix proteins (P_adj_ = 3.5 × 10_−25_) (such as genes encoding for collagens (P_adj_ = 0.4 × 10_−10_)), and the purple module was enriched for genes involved in skeletal system development (P_adj_ = 2.3 × 10_−10_) and osteoblast differentiation (P_adj_ = 2.0 × 10_−6_) (**Figure 1 and Supplemental File 2**). Given that network modules represented distinct biological processes, including those involved in mineralization and osteoblast activity, we were confident it would provide a platform for identifying core genes related to mineralization that potentially underlie BMD GWAS associations.

### II. Identification of co-expression modules enriched for genes implicated by GWAS

To identify modules of co-expressed genes informative for GWAS, we first determined if any of the 65 modules were enriched for genes that overlapped GWAS associations. Using data from the two largest GWASs performed at the time, one study of Dual Energy X-Absorptiometry (DEXA) derived areal BMD measures at the lumbar spine and femoral neck ^8^ (“Estrada *et al.* GWAS”; N=32,961) and one study of ultrasound determined heel estimated BMD (eBMD) ^9^ (“Kemp *et al.* GWAS”, N=142,487), we developed a list of 789 human genes (N_Estrada_ = 179, N_Kemp_ = 701, (91 shared genes)) intersecting BMD GWAS loci. A total of 723 (92%) of these had mouse homologs in the network (**Supplemental File 3 and 4**). Of the 65 modules in the network, 13 were enriched for mouse homologs of human genes implicated by BMD GWAS (Fisher’s exact test, P_adj_ < 0.05) (**Supplemental File 5 and Figure 2A**). Additionally, we performed stratified LD score regression by calculating the BMD heritability partitioned by SNPs surrounding genes in each module using the Kemp *et al.* GWAS ^9, 25^. We found 16 modules enriched for partitioned BMD heritability, including nine of the 13 enriched for BMD GWAS implicated genes (**Figure 2B and Supplemental File 6**).

**Figure 2.**
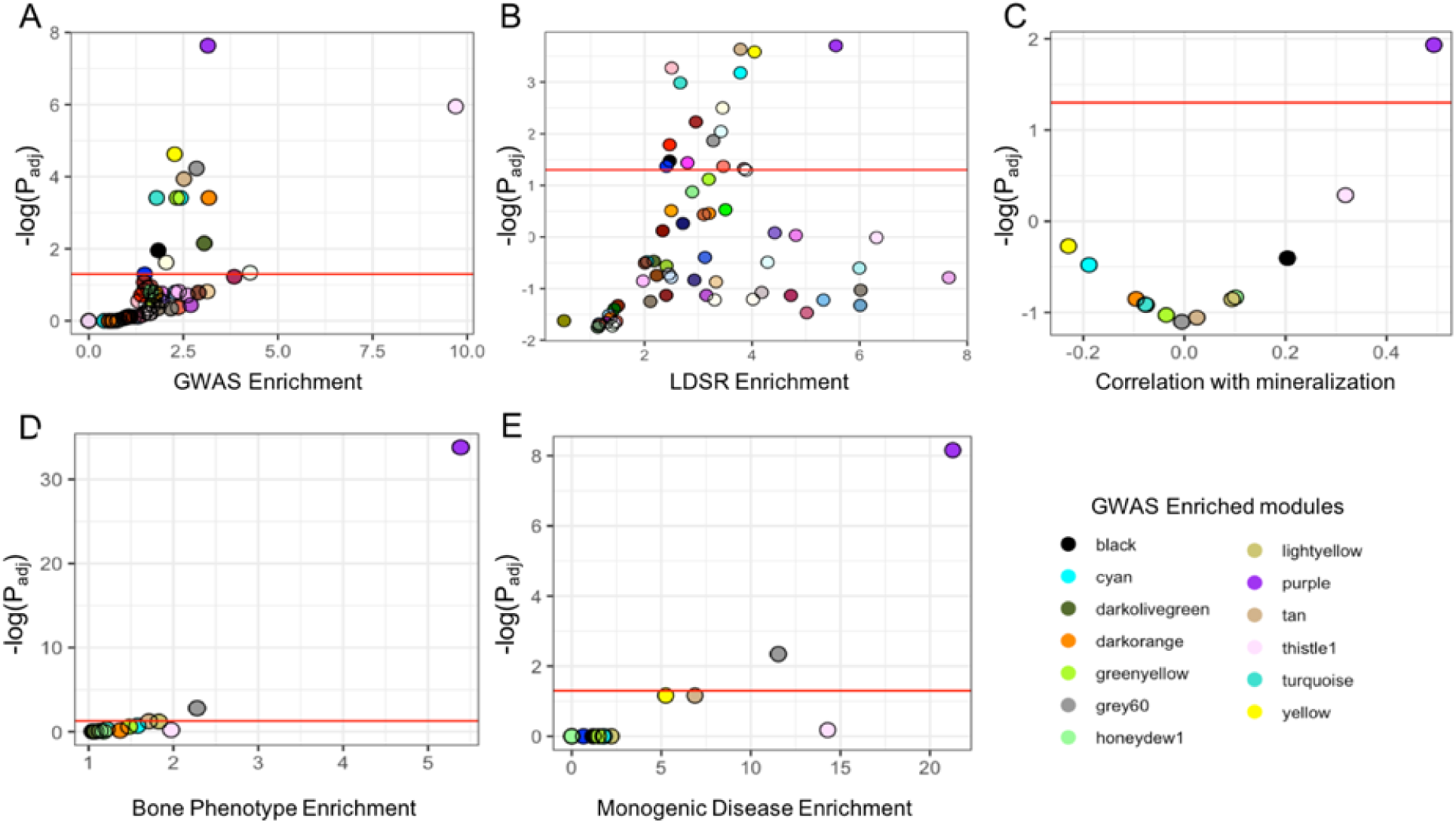
The purple module is enriched for genes with core-like properties. (A) Module enrichments for genes overlapping a BMD GWAS association. (B) Enrichments for partitioned BMD heritability for each module determined using stratified LD score regression. (C) Correlation between each module eigengene and *in vitro* mineralization. (D) Module enrichments for genes that, when knocked out, produced a bone phenotype and (E) human monogenic bone disease genes. Red line in each panel represents Padj < 0.05.

### III. The purple module is enriched for genes with core-like properties

Next, we focused on identifying which of the 13 modules identified above were enriched for genes with core-like properties. To accomplish this, we selected modules using biologically motivated criteria which likely reflected the properties of core genes. First, we compared the 13 module eigengenes with *in vitro* mineralization using osteogenic cultures from the same 42 CC strains used in the construction of the co-expression network (**Supplemental Figure 1**). Only one, the purple module, had a pattern of expression that was significantly correlated with mineralization (*r =* 0.49, P_adj_ = 0.012), suggesting the purple module was enriched for genes with a direct role in mineralization (**Figure 2C and Supplemental Figure 2**).

Core genes have been broadly defined as those that directly influence a disease-relevant biological processes ^11, 12^. Thus, severe perturbation of a core gene is more likely to result in a significant impact on a phenotype, as in the case of a mouse knockout or human monogenic disease. We identified all gene knockouts that produced a bone phenotype, defined as either a change in BMD, bone mineral content (BMC), abnormal bone morphology, or abnormal bone cell activity, by utilizing mouse knockout phenotype data from several databases ^26–29^ (**Supplemental File 7**). Of the 13 modules enriched for BMD GWAS genes, two were enriched for genes whose deficiency impacted bone in mice (**Figure 2D**). The purple module was the most significantly enriched (OR=5.4, P_adj_ = 1.6 × 10_−34_). We also compiled a list of 35 known drivers of monogenic bone diseases associated with osteoblast dysfunction, including osteogenesis imperfecta, hyperostosis, and osteosclerosis (**Supplemental File 8**) ^30–34^. Again, the purple module, containing 11 of 35 (31.4%) monogenic disease genes, was the most significantly enriched (OR = 21.3, Padj = 6.9 × 10_−9_) (**Figure 2E**). Together, these independent lines of evidence suggested the purple module was enriched for genes with core-like properties.

### IV. New BMD GWAS associations further support the purple module as a core gene module

While we were analyzing the Kemp *et al.* GWAS data, an extension of this study, with a significantly increased eBMD sample size, was published (“Morris *et al.* GWAS”) ^14^. The Estrada *et al.* (N=32,961) and Kemp *et al.* (N=142,487) GWASs identified 56 and 307 conditionally independent associations, respectively ^8, 9^. In comparison, the Morris *et al.* GWAS (N=426,824) identified 1103 eBMD associations; an increase of over 3.5-fold ^14^. The associations identified in the Morris *et al.* GWAS overlapped 1581 genes, as compared to 789 in the Estrada *et al.* and Kemp *et al.* GWASs (**Supplemental File 9**). Assuming the genetic architecture of BMD is consistent with the omnigenic model, we expected the inclusion of the Morris *et al.* GWAS data to increase the number of modules enriched for GWAS implicated genes. Consistent with this hypothesis, the number of modules enriched for GWAS-implicated genes doubled (N_Kemp_ = 13, N_Morris_ = 26) using the Morris *et al.* GWAS (**Figure 3A**) (**Supplemental File 10**). As observed in the first analysis, most (18/26, 69%) of the new modules enriched for GWAS-implicated genes were also enriched for partitioned BMD heritability (**Supplemental File 11 and Figure 3C**). These new modules were enriched for genes involved in general biological processes such as RNA splicing (brown module, Padj = 4.0 × 10_−11_), cell junctions (floralwhite module, Padj = 6.2 × 10_−3_), cell motor activity (orange, Padj = 6.6 × 10_−3_), the cell cycle (lightgreen, Padj = 3.2 × 10_−4_), ER to Golgi trafficking (salmon, Padj = 1.8 × 10_−2_), and the glycolytic process (red, Padj = 1.1 × 10_−13_), and not processes specific to osteoblast activity and/or mineralization **(Supplemental File 2)**.

**Figure 3.**
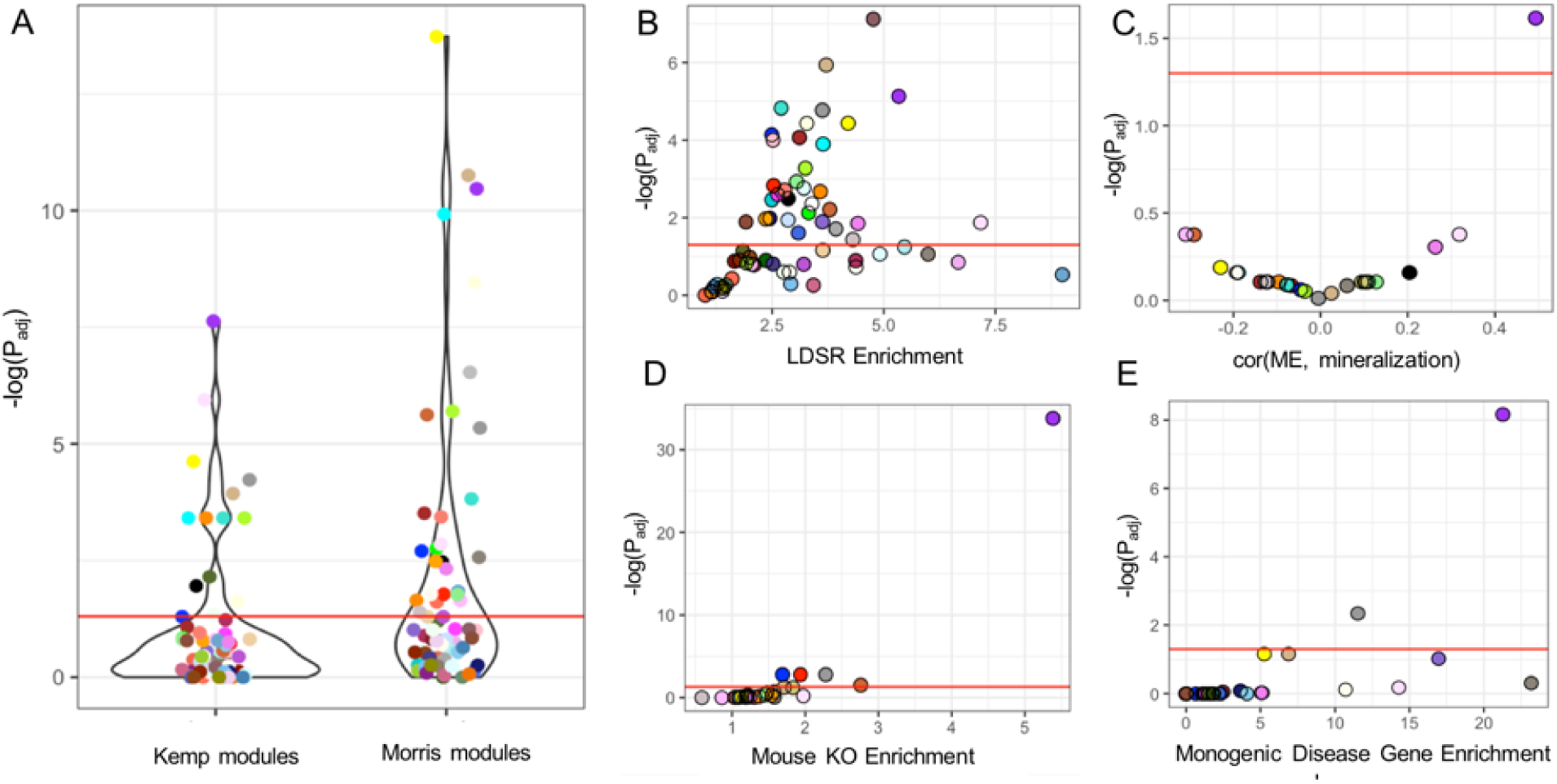
The purple module was the only core module after increasing the number of analyzed GWAS associations by 3.5-fold. (A) A greater number of modules were identified as enriched for GWAS implicated genes in the Morris *et al.* GWAS versus the Kemp *et al.* GWAS. (B) Module enrichments for partitioned BMD heritability for each module determined using stratified LD score regression. (C) Correlation between each module eigengene and *in vitro* mineralization. (D) Module enrichments for genes that, when knocked out, produced a bone phenotype and (E) human monogenic bone disease genes. Red line in each panel represents Padj < 0.05.

Similar to the analysis of the Kemp *et al.* data, the purple module was among the most enriched for GWAS implicated genes (OR = 2.67, Padj = 3.4 × 10_−11_) (**Figures 3A**) and BMD heritability captured (OR = 5.8, Padj = 4.7 × 10_−6_) (**Figures 3B**). Using the Estrada *et al.* and Kemp *et al.* GWAS, the purple module contained 45 genes implicated by GWAS (OR = 3.15, Padj = 2.3 × 10_−8_) (5.7% of GWAS-implicated genes; 8.9% of purple module genes) and explained 27% of the SNP-heritability (h_g2_) in the study, or 4.6% of the total heritability. Using the Morris *et al.* GWAS, the number of purple module genes implicated by GWAS increased to 77 (OR = 2.7, Padj = 3.4 × 10_11_) (4.9% of GWAS-implicated genes; 15.2% of purple module genes) explaining 25.3% of the h_g2_, or 5.4% of the total heritability. Additionally, the purple module was still the only one correlated with *in vitro* mineralization (**Figure 3D**), the most significantly enriched for genes eliciting a bone phenotype when knocked-out in mice (**Figure 3E**), and human monogenic bone disease genes (**Figure 3F**). These data indicate that even with a significant increase in the number of GWAS-implicated genes included in the analysis, the purple module is the only one enriched for genes with core like properties.

### V. The purple module contains genes belonging to one of two distinct transcriptional programs across osteoblast differentiation

The purple module was enriched for GO categories important for the function of osteoblasts. Consistent with this observation, it contained many genes known to play key roles in osteoblast differentiation and mineralization, including *Runx2* ^13^, *Sp7* ^35^, *Sost* ^36, 37^, *Bglap* ^38^, *Alpl* ^39^, among many others (**Supplemental File 12**). Thus, to further investigate the purple module, we evaluated the expression of its genes with regards to osteoblast differentiation. To do this, we utilized transcriptomic profiles collected from purified osteoblasts at multiple time points across differentiation (GSE54461). Using k-means clustering, we found that the genes within the purple module clearly partitioned into two distinct transcriptional profiles with regards to differentiation (**Figure 4A,B**). We have termed these groups the Early Differentiation Submodule (EDS; high expression early and low expression late) (N=192 transcripts; 175 unique genes) and the Late Differentiation Submodule (LDS; low expression early and high expression late) (N=423 transcripts; 323 unique genes).

**Figure 4.**
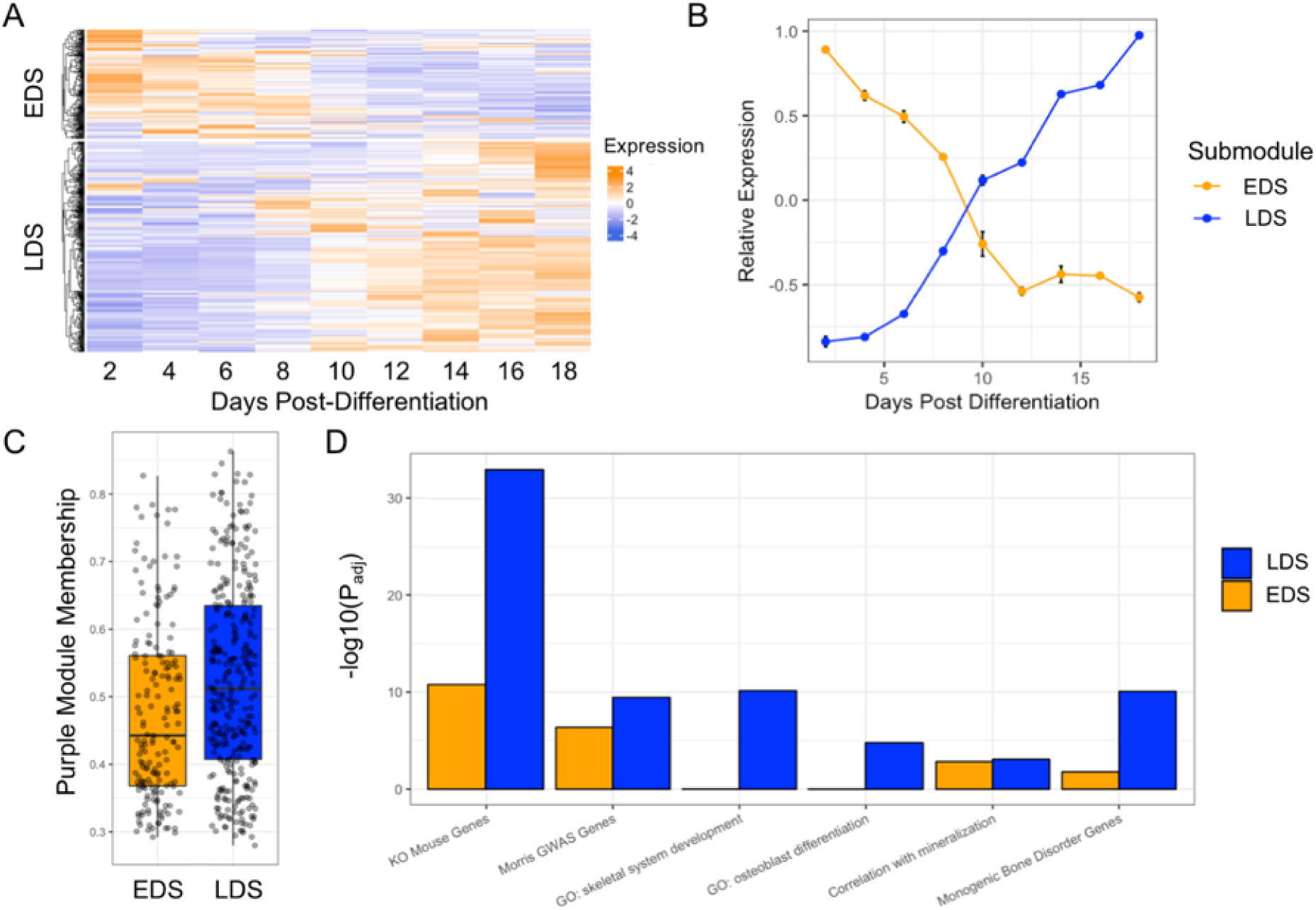
The purple module consists of genes representing two distinct transcriptional profiles across osteoblast differentiation, one of which, the late differentiation submodule (LDS), is more enriched for genes with properties consistent with core genes for mineralization. (A) Purple module genes show two distinct patterns of expression across differentiation, (B) Genes in cluster 1 (or the early differentiation submodule; EDS; N=175 genes) are expressed high early in osteoblast differentiation. Genes in cluster 2 (or the late differentiation submodule; LDS; N=323 genes) are expressed high late in osteoblast differentiation. (C) LDS genes have a significantly higher purple module membership score (P = 3.0 × 10_−4_). (D) The LDS is more significantly enriched than the EDS for genes implicated by BMD GWAS in humans, associated with GO terms for bone development, for genes that when knocked out, produce a bone phenotype, and for genes involved in monogenic bone disorders.

We assessed whether there were differences between the EDS and the LDS with regard to network parameters and their enrichment for functional annotations seen in the purple module. We first looked at intramodular connectivity, measured by module membership (correlation between the expression of each gene and the module eigengene). On average, LDS genes had higher module membership scores than EDS genes (P = 3.0 × 10_−4_) (**Figure 4C**), suggesting they may play more critical roles in the context of overall module behavior. Additionally, the LDS was more significantly enriched for genes implicated by GWAS (OR = 3.0, Padj = 5.2 × 10_−10_), osteoblast relevant GO terms (e.g. “ossification” (Padj = 3.24×10_−14_), skeletal development” (Padj = 9.6 × 10_−11_), “osteoblast differentiation” (Padj = 1.4 × 10_−4_), and “biomineral tissue development” (Padj = 4.1×10_−6_), genes that when knocked-out result in a bone phenotype (OR = 7.3, Padj = 1.1 × 10_−33_) and monogenic bone disease genes (OR = 33.2, Padj = 8.4 × 10_−11_) (Figure 4D). As one would expect based on their higher expression later in differentiation, many of the most well-known regulators of mineralization, such as *Phospho1* ^40^, *Bglap* ^41^, *Fam20c* ^42^, *Mepe* ^43^, *Phex* ^44^, to name a few, were members of the LDS (**Supplemental File 12**). These observations, together with the fact that LDS genes are expressed at high levels during late differentiation, coincident with when the osteoblasts are actively mineralizing, suggest that LDS contains genes with core-like properties specific for the process of mineralization. For all downstream analyses we focused on the LDS.

### VI. BMD-associated variants in GWAS loci harboring LDS genes overlap active regulatory elements in osteoblasts

Based on the fact that the LDS is enriched for genes involved in osteoblast differentiation and that mineralization is fundamental in the regulation of BMD, we anticipate that many of the genes in the LDS are true core genes and causal genetic drivers of BMD. If true, then BMD-associated variants in associations harboring LDS genes should regulate the expression of LDS genes in osteoblasts. To test this, we utilized histone modification data from the Roadmap Epigenome Project ^45^. In the Morris *et al.* BMD GWAS, 48 LDS genes overlapped 84 associations (7.6% of the 1103 total; a subset of LDS genes overlapped multiple clustered associations). For each of the 84 independently associated lead (i.e., most significant) SNPs, we analyzed histone modifications across the osteoblast genome and observed that they were more likely to overlap regions marked by modifications associated with active regulatory elements such as H3K4me1 (2.8x enrichment, P < 1 × 10_−3_), H3Kme2 (3.2x enrichment, P < 1 × 10_−3_), H3K4me3 (3.8x enrichment, P < 1 × 10_−3_), and H3K27ac (2.6x enrichment, P < 1 × 10_−3_) relative to 1000 sets of random SNPs matched for allele frequency and distance from a transcription start site (**Figure 5**). Additionally, we observed depletion of LDS SNPs in heterochromatic regions, marked by H3K9me3 (0.14x depletion, P < 1 × 10_−3_).

**Figure 5.**
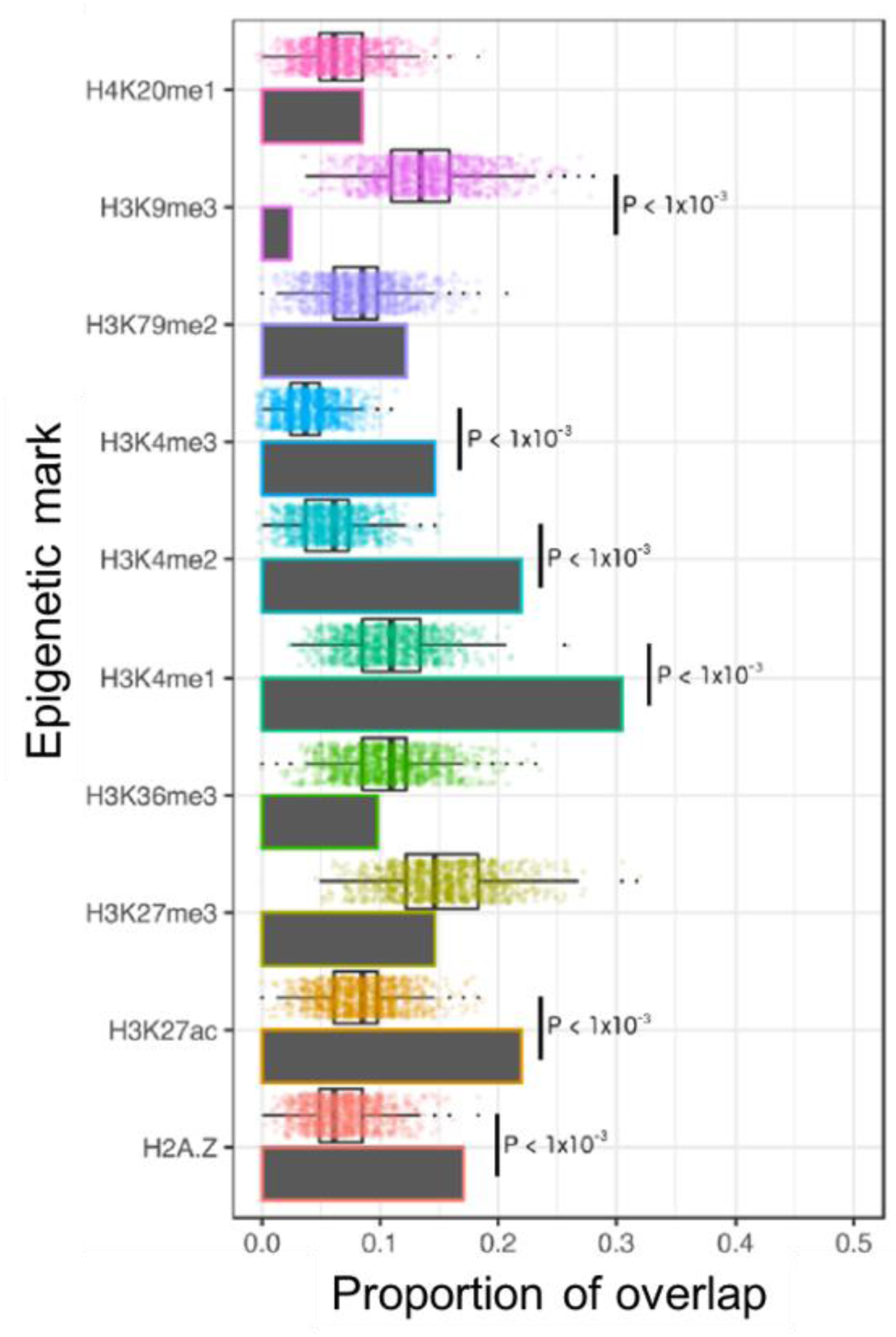
Lead SNPs for GWAS associations harboring LDS genes overlap active regulatory elements in osteoblasts. Grey bars represent the proportion of LDS SNPs (n = 84) that overlap each of the epigenetic marks measured in osteoblasts. Box and dot plots represent the proportion of each set of random SNPs (N = 1000) (matched to the LDS SNPs for MAF and distance from TSS) overlapping each epigenetic mark measured in osteoblasts.

To determine if the enrichments were specific to osteoblasts, we calculated the ratio between the LDS BMD set overlap and the mean random set overlap across all 129 Roadmap tissues and cell-types. For all activating marks (H3K27ac, H3K4me1, H3K4me2, H3K4me3) osteoblasts were in the top 10% when tissues were ranked based on the overlap ratio (**Supplemental File 13**). The tissues for which the random sets had a higher ratio included cell types within the same lineage as osteoblasts, such as mesenchymal stem cell (MSC) derived chondrocytes and other MSC-derived tissues including adipose and skeletal muscle. These data support the premise that loci harboring LDS genes impact BMD through the regulation of gene expression in osteoblasts, further supporting the causality of LDS genes.

### VII. LDS genes *CADM1*, *B4GALNT3*, *DOCK9*, and *GPR133* are novel genetic determinants of BMD

The overarching goal of this study was to identify causal genes from a module enriched for genes with core-like properties underlying BMD GWAS associations. As described above, 48 (14.9%) LDS genes overlapped an eBMD GWAS association from the Morris *et al.* study. To further identify those with strong evidence of being causal, we utilized expression quantitative trait locus (eQTL) data from the Gene Tissue Expression (GTEx) project to identify local eQTL colocalizing with BMD associations ^46^. We also used total body BMD data on LDS gene knockouts collected as part of the International Mouse Phenotyping Consortium (IMPC) ^27^. Together, these data allowed us to directly link BMD associated variants to LDS genes and LDS genes to pathways regulating BMD. We performed a colocalization analysis for each eQTL/BMD association pair for all 48 genes in 48 GTEx tissues and identified 12 LDS genes with colocalizing eQTL (PP4>0.7) (**Supplemental File 14 and Figures 6A, B, C, and D**). We also queried each of 12 LDS genes with a colocalizing eQTL and found that BMD had been measured by the IMPC in 5 mutants. Of these, four genes (*Cadm1*, *B4galnt3*, *Dock9*, and *Adgrd1*) had significantly altered total body BMD (Padj < 0.05) (**Supplemental File 15 and Figures 6E, F, G and H**). For *Cadm1* and *Dock9* the direction of effect inferred from the eQTL/BMD association matched the direction of the effect observed in the mouse knockout; however, for *B4galnt3* and *Adgrd1* the directions did not match (**Supplemental File 15**).

**Figure 6.**
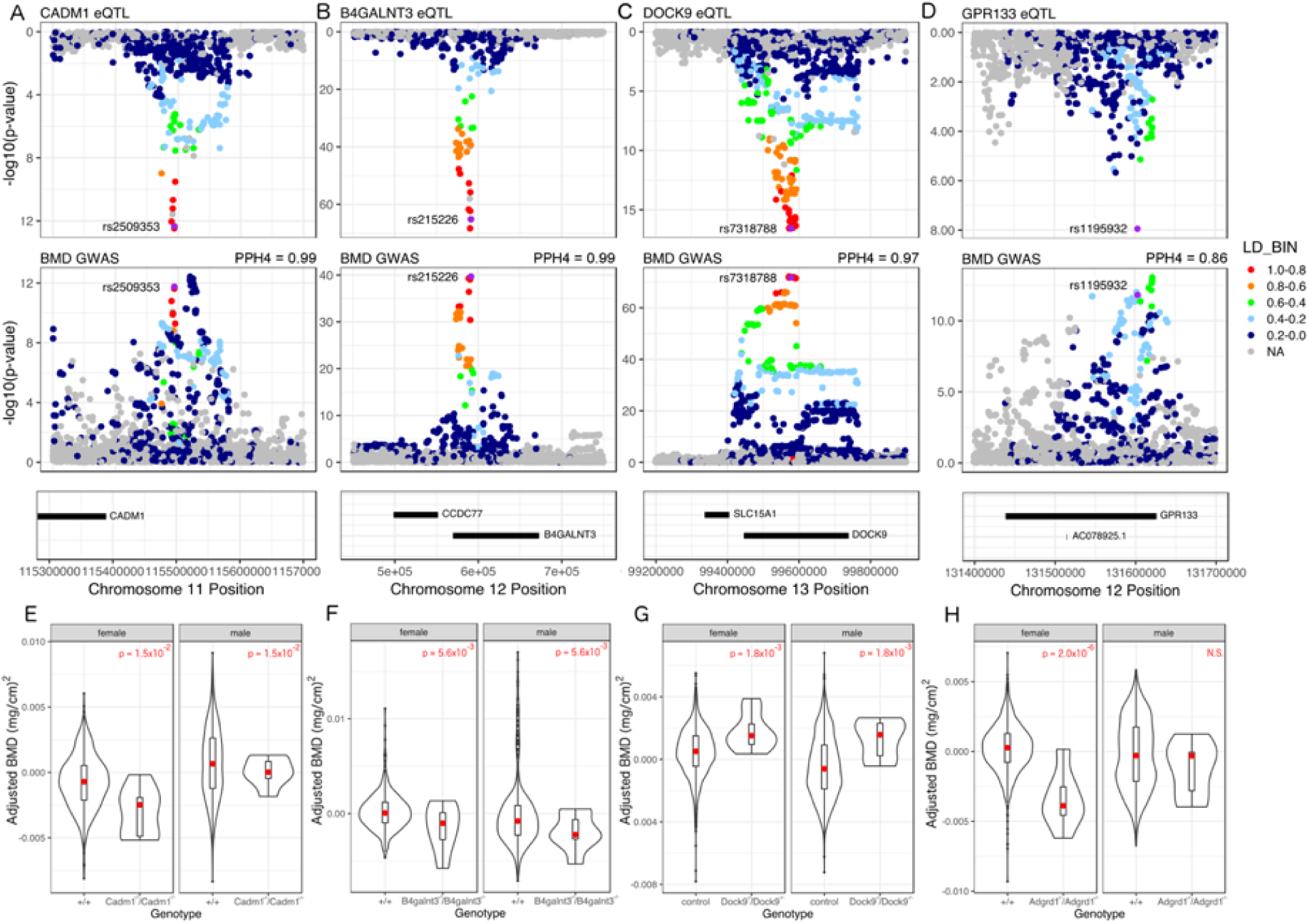
CADM1, B4GALNT3, DOCK9, and ADGRD1 (aka GPR133) are novel genetic regulators of BMD. (A-D) All four genes have an eQTL in at least one tissue in the GTEx database that colocalizes with a proximal BMD GWAS association. (E-H) Knockout mice from the KOMP for each gene exhibit altered BMD.

Lastly, we evaluated network parameters of *Cadm1*, *B4galnt3*, *Dock9* and *Adgrd1*. We observed that *Cadm1* and *B4galnt3* were ranked in the top 20 based on LDS connectivity (**Supplemental File 12**). In fact, *Cadm1* was the 2nd most highly connected gene. Together, the four genes had, on average, higher module membership than the average LDS gene (0.72 vs. 0.52; P = 0.002). In support of the importance of connectivity in the LDS, we observed that more highly connected LDS genes were more likely (P=0.008) to overlap a BMD GWAS locus (**Supplemental Figure 3A**) and there was a strong positive correlation between connectivity and *in vitro* mineralization for all LDS genes (r = 0.71, P< 2.2 × 10_−16_) (**Supplemental Figure 3B**). These data suggest that connectivity is an important feature of the LDS and a strong proxy for biological importance. Furthermore, these data strongly support *CADM1*, *B4GALNT3*, *DOCK9* and *ADGRD1* as genetic drivers of BMD in humans.

## Discussion

Osteoporosis is an increasingly common disease associated with reduced BMD and negative health outcomes, namely fracture 1. Despite its significant genetic component, we do not fully understand the genes and mechanisms that influence osteoporosis and its determinants, such as BMD. Moreover, current therapeutics for osteoporosis have been associated with rare side effects, leading to decreased compliance ^47^. Identification of the causal genes with core-like properties that regulate BMD will help us to further understand the etiology of osteoporosis and lead to the development of novel therapeutics. In this study, we identified the LDS, a co-expression submodule enriched for genes with core-like properties influencing BMD, by integrating a cell- and timepoint-specific co-expression network with the results of BMD GWAS. We then used this information to identify four LDS genes that overlapped GWAS loci, had colocalizing eQTL, and altered BMD in knockouts, suggesting they are causal for their respective BMD GWAS association.

In this work, we hypothesized that the genes underlying BMD GWAS associations are not created equal with respect to “biological importance” or membership in pathways with direct impacts on bone mass. Substantial prior evidence supports this prediction ^48, 49^ and it is one of the primary tenets of the omnigenic model ^11, 12^. Identification of genes whose activity or function is more proximal to BMD is important for a number of reasons. First, the identification of genes with core-like properties has the potential to identify critical new players in pathways known to directly impact bone and to uncover new processes essential to skeletal growth and maintenance. Second, it provides a way to prioritize hundreds of BMD GWAS loci for further investigation. Third, based on their central role in the regulation of BMD, it is logical to use the concept of a core gene as a way to prioritize gene discovery in the context of selecting promising new therapeutic targets for evaluation.

The omnigenic model uses a strict statistical definition to define core genes and many have debated the utility of this designation^11, 15–17^. Some have argued that focusing on core genes underestimates the complexity of complex traits, attributing biologically nuanced diseases to a small set of genes ^13^. Others have argued that the focus should not be on thoroughly defining core genes, but instead on identifying the underlying biological pathways and mechanisms ^14^. In practice, it is likely that the designation of core genes follows a spectrum rather than a discrete classification. If so, then it should be possible to rank genes based on their continuous “core” attributes, which would be analogous to ranking genes based on their proximity to a disease or phenotype. In essence, that is what we have done in the current study with the goal of identifying genes on the end of the “core” attribute distribution for mineralization. Importantly, it is not likely that all genes in the LDS are causal genetic drivers or, if they are causal, it is possible that several will have few core attributes. However, based on our analysis and results, it is likely that many are causal genes that participate in “core” pathways and processes that directly impact mineralization, bone formation, and BMD.

As we have previously demonstrated ^50, 51^, there are a number of advantages to using co-expression networks to inform GWAS. First, it allowed us to group genes across the genome based on function and pathway membership and then identify groups of functionally similar genes that had core-like properties. Second, it allowed us to predict the function of potentially casual genetic drivers of BMD. Based on the strong GO enrichments and membership of genes with well-known roles in bone formation and mineralization, it is likely that all LDS genes, including those with no known function, impact mineralization in some manner. The idea of the LDS playing a central role in bone formation was further supported by the strong overlap observed between lead BMD GWAS SNPs for associations containing LDS genes regulatory elements in osteoblasts. Third, it begins to provide a systems-level context for causal genetic drivers. Once genes underlying GWAS loci are identified it is then important to begin to understand their role in complicated cellular networks, defining how a set of genetic variants may converge on multiple genes all involved in a particular process. We can use the LDS to begin to identify sets of variants that all work to influence genes which impact mineralization and the hierarchy of relationships between these genes.

This work extends our use of co-expression networks to inform GWAS. Previously, we used a network generated using cortical bone expression profiles from the Hybrid Mouse Diversity Panel to identify two “osteoblast” modules (enriched for genes involved in osteoblast differentiation and function) enriched for genes implicated by BMD GWAS. We used these modules to identify 35 genes potentially causal for GWAS loci, including two (*MARK3* and *SPTBN1*) that we experimentally validated their involvement in BMD. Comparing the two modules to the LDS we observed a modest overlap (96 of 323 genes; 29.7%), even though they both demonstrated a strong “osteoblast” enrichment signature. While a number of differences (microarray vs. RNA-seq transcriptomic data, different mouse populations, etc.) confound the interpretation of the seemingly low overlap, it is likely due in large part to our use of osteoblast-specific network capturing the transcriptome at peak mineralization instead of the whole bone tissue representing a small number of osteoblasts.

We provided strong supporting evidence that four LDS genes (*CADM1*, *B4GALNT3*, *DOCK9* and *GPR133*) are novel regulators of BMD and causal for their respective GWAS association. Prior to this study, none of these genes had been directly connected to the regulation of BMD. *CADM1* (Cell Adhesion Molecule 1) is a ubiquitously expressed cell adhesion molecule involved in many biological processes, including cancer, spermatogenesis, and neuronal/mast/epithelial cell function^52–54^ that had been implicated in osteoclast proliferation and activity^55^ and as an osteoblast-specific marker in the context of osteosarcoma^56, 57^. *B4GALNT3* (Beta-1,4-N-Acetyl-Galactosaminyltransferase 3) is a glycosyltransferase that transfers N-acetylgalactosaine (GalNAc) onto glucosyl residues, thus forming N,N-prime-diacetyllactoseadiamine (LacdiNAc), which serves as a terminal structure of cell surface N-glycans that contributes to cell signaling ^51, 52^. *B4GALNT3* is expressed in bone and associated with circulating levels of sclerostin ^53–55^. *DOCK9* (Dedicator of Cytokinesis 9) is a guanine nucleotide-exchange factor (GEF) that activates Cdc42^58^, which has been shown to regulate osteoclast differentiation and ossification ^57, 58^. *GPR133* (Adhesion G Protein-Coupled Receptor D1) is a G protein-coupled receptor that participates in cell-cell and cell-matrix interactions ^59^. Our results demonstrate the utility of the LDS in broadening our understanding of the molecular and genetic basis of BMD.

Our study is not without limitations. First, we used gene expression data from the mouse as a discovery platform, however this may limit the translational applications of the work due biological differences and missing homologs between mouse and human. Secondly, this was not a comprehensive study of the genetic effects driving osteoporosis, because we focused exclusively on the contributions of just one cell type, bone-forming osteoblasts. In future work, this approach could also be applied to other bone cell types. For example, one could use *in vitro* measures of osteoclast activity as a filter to identify groups of genes influencing bone resorption, and ultimately BMD. Finally, the eQTL comparisons made in this study were not derived from expression data in bone tissue, as bone tissue expression was not measured in the GTEx project. While we identified colocalizing eQTL in other tissues, these eQTL may be irrelevant to BMD or the direction of eQTL effects in non-bone tissues may not reflect the direction of effect in osteoblasts.

While we identified four novel regulators of bone mineral density, there is still much to be gleaned from the late differentiation submodule. The LDS is a promising resource for two key applications: (1) causal gene discovery and functional follow up and (2) studying the impact of genetic variation on biological networks and complex phenotypes. There are still many genes with no known connection to BMD in the LDS that are likely important to osteoblast biology and mineralization. Additionally, the LDS is not just a list of candidate genes; it also provides insight into the molecular hierarchy driving osteoblast differentiation and mineralization, which can demonstrate how genetic variation impacts biological networks. The network topology of the LDS can also be used to infer the causal relationships between genetic variants and the many genes that influence osteoblast activity. Moving forward, the LDS can serve as a platform for the identification of novel determinants of BMD and for furthering our understanding of the nuanced relationship between genetic variation, molecular phenotypes, and complex traits.

In summary, we have used an integrative, network-based method to identify core genes for the process of mineralization and BMD. While the definition of a core gene is still open to debate, we found the expected properties of core genes are effective lenses through which to contextualize GWAS associations. Integrating gene co-expression networks, GWAS data, *in vitro* and *in vivo* phenotypic data reflecting “core” properties, and eQTL information has led us to a more complete understanding of the biology and genetics of BMD.

## Methods

### RNA-seq

Neonatal collaborative cross heads were received from the University of North Carolina. At UNC, neonatal (3-5 days) collaborative cross mice were euthanized by CO_2_, decapitated onto paper towels soaked in 70% ethanol, and placed in cold PBS on ice for overnight shipping. Once received, calvaria were dissected, paying special attention to brain and interparietal bone removal. Isolated calvaria were placed in 24 well plates containing 0.5 mL of digest solution (0.05% trypsin and 1.5 U/ml collagenase P) and incubated on a rocking platform at 37 degrees during six, fifteen-minute digestions in 0.5 mL of digestion solution. Fraction 1 is discarded and fractions 2-6 are collected. Fractions 2-6 are added to an equal volume of cold plating media (89 mL DMEM, 1 mL 100x Pen/Strep solution, and 10 mL Lot tested FBS). The resulting cells are filtered using a 70-100 mm cell strainer to remove clots, centrifuged at 1000 rpm for 5 minutes and re-suspended in 0.5 ml plating media. The resulting cells are plated in a T25 flask. 24 hours later, cells are washed with PBS, treated with trypsin, counted, and plated at a density of 1.5×10_5_ cells per well in a 12-well plate, and allowed to grow to confluence for 48 hours. After 48 hours of growth, cells are switched to differentiation media (10 mL lot tested FCS, 1 mL 100x Pen/Strep solution, 283.8 uL ascorbic acid (0.1 M), 400 uL B-glycerol phosphate (1 M), and 88.3 mL alpha-MEM per 100 mL) and allowed to differentiate for 10 days. On day 10, total RNA was extracted from the mineralized cultures using *mir*Vana RNA isolation kit (ThermoFisher Scientific).

RNA-Seq libraries were constructed from 200 ng of total RNA using Illumina TruSeq Stranded Total RNA with Ribo-Zero Gold sample prep kits (Illumina, Carlsbad, CA). Constructed libraries contained RNAs >200 nt (both unpolyadenylated and polyadenylated) and were depleted of cytoplasmic and mitochondrial rRNAs. An average of 39.7 million 2 x 75 bp paired-end reads were generated for each sample on an Illumina NextSeq 500 (Illumina, Carlsbad, CA). FastQC was used to evaluate the quality of the reads, and all samples passed the QC stage ^59^. Reads were mapped to the eight collaborative cross founder transcriptomes based on build mm9 using Bowtie, and quantified using EMASE ^60^. EMASE output transcript level expression estimates calculated by assigning multi-mapping reads across the genome using and expectation-maximization algorithm to allocate reads that differentiate between genes, then isoforms of a gene, and then alleles.

### WGCNA network construction

Estimated transcript count data was used as the basis for co-expression network construction. We removed transcripts with less than an average tpm <= 0.3 tpm across all samples, resulting in 29,000 transcripts used to construct the network. We used a variance stabilizing transformation from the DESeq2 package that decouples the variance from the mean ^61^. Next, we used PEER in order to remove latent confounding batch effects from our data ^62^. As per PEER recommendations, we estimated PEER factors equal to one quarter of the number of samples (N = 24) and included covariates in the calculation. We carried out the downstream analysis with the residual values from PEER transformation. Finally, we used quantile normalization to match the distribution of each of the samples in the analysis.

The resulting expression data was used to construct a signed, weighted gene co-expression network using the weighted gene co-expression network analysis (WGCNA) package ^63^. There were no evident outliers from the hierarchical clustering analysis. The pickSoftThreshold() function from the wgcna package was used to determine the power used to calculate the network. The minimum power value that had an R_2_ >= 0.9 for the scale-free topology model fit was used, and the network was calculated using a power of 9. We then used the blockwiseModules() function to construct a signed network with a merge cut height of 0.15, and a minimum module size of 20 genes. Using WGCNA, we constructed a signed network composed of 65 modules of co-expressed genes, with an average of 292 genes per module.

### Gene Ontology Analysis

For those modules that were enriched for BMD GWAS genes, we conducted gene ontology analysis to identify the functional categories represented by each module. Using the ToppFun tool on the ToppGene site, we identified the significantly enriched categories for GO molecular functions, GO biological processes, GO cellular components, human and mouse phenotypes, and pathways ^64^. The significance cutoffs reported for these enrichments were Benjamini & Hochberg corrected FDR q-values.

### Creating BMD GWAS list

In order to identify co-expression modules enriched for BMD GWAS genes, we identified all genes overlapping a BMD GWAS locus using the 2012 and 2017 BMD GWAS ^8, 9^. For each BMD locus, a bin was defined by the furthest upstream and downstream SNPs with LD >= 0.7 as calculated from the European populations in the 1000 genomes phase III data identified using the LDLink LDProxy tool ^65^. Then, using the Genomic Ranges tool, we identified all genes from the GRCh37/hg19 Ensembl gene set overlapping a BMD GWAS bin ^66, 67^. If not gene intersected a bin, we identified the nearest upstream and downstream genes from the bin. The Estrada GWAS resulted in 179 genes and the eBMD GWAS resulted in 701 genes, resulting in a list of 731 unique genes. We converted the list of human genes to mouse homologs.

### BMD GWAS gene enrichment

In order to identify modules of genes enriched for GWAS genes, we used a fisher’s exact test to measure the statistical significance of the representation of GWAS genes in each module. We then applied a Bonferroni correction to correct for testing the enrichment of all 65 modules, and applied a significance cutoff of 0.05 to the adjusted p-values, resulting in 13 modules of genes enriched for 2012 and 2017 GWAS genes, and 26 modules of genes enriched for 2012, 2017, and 2018 GWAS genes.

### LD Score Regression

In order to evaluate the relevance of the BMD GWAS gene enriched modules we calculated the partitioned heritability of the SNPs in the regions surrounding the genes in each module. We used the LD score regression method, which takes gene lists as an input and returns the enrichment of the associated SNP set for heritability for the tested trait. For each set of modules we tested using this method, we corrected the enrichment p-values for multiple testing using a Bonferroni correction, and applied a p-value cutoff of 0.05 to the adjusted p-values.

### *In vitro* mineralization measurement and correlation

In order to identify the modules of coexpressed genes with patterns of expression correlated with mineralization, we measured *in vitro* mineralization in osteogenic cells from the calvaria of 42 strains of collaborative cross mice. After 10 days of differentiation and mineral production, cells are washed with PBS and treated with 10% NBF (1 mL per well) and incubated at room temperature for 15 minutes. The NBF is removed and cells are washed with H_2_O (1mL x 2). Next, wells are stained with alizarin red (0.5 mLs, 40 mM @ pH 5.6) for 20 minutes on a shake plate at 120 rpm. Alizarin red stain is then removed, and cells are washed 5 times with deionized H_2_O for 5 minutes on a shake plate at 180 rpm. Once rinsed, the mineralized wells are scanned, and .tiff images are retained to extract geometric parameters of the mineral deposits. After imaging, the wells are de-stained by incubation with 5% perchloric acid (1 mL) at room temperature for 5 minutes while shaking at 120 rpm. Eluent is collected and read at 405 nm. The levels of *in vitro* mineralization varied significantly across the population, with a 63-fold change from the highest to lowest mineralization samples (max_mmAR = 2.995993, min_mmAR = 0.04719).

In this population, *in vitro* mineralization had a heritability of 47.8%(p=1.8×10_−46_), indicating that the between-strain variation is larger than the within strain variation and that there is a genetic contribution to the process of mineralization. Using the WGCNA package, the eigengene of each module was calculated, and the correlation between the eigengene and the in vitro mineralization phenotype was calculated using the cor() function in R. The p-values associated with the correlation between the module eigengenes and *in vitro* mineralization were corrected for multiple testing using a Bonferroni correction and a p-value cutoff of 0.05 was applied to the adjusted p-values.

### Module enrichment for genes with associated bone phenotypes and monogenic bone disease

In order to identify modules of coexpressed BMD GWAS genes that are enriched for genes with bone phenotype annotations, we curated a list of genes which produce a bone phenotype when knocked out. We used four databases of gene perturbations that result in bone phenotypes, including genes annotated with a bone phenotype in the Mouse Genome Informatics database (MGI), the Origins of Bone and Cartilage Disease (OBCD) database, the International Mouse Phenotyping Consortium (IMPC), and the Bonebase Database ^26–29^. Specifically, we pulled BMD, altered bone morphology, altered bone cell activity, changes in ossification or mineralization, or association with a known bone disease from the MGI database. The OBCD database contained genes with changes in bone mineral content (BMC), bone volume fraction (BV/TV), and BMD of the femur and BMD of the vertebra. We mined the IMPC database for any genes with altered BMD, and we pulled all Bonebase genes with altered BV/TV in the femur or vertebra. This resulted in a list of 923 unique “bone” genes (**Supplemental File 7**).

We also curated a list of genes associated with monogenic bone disorders using a literature review, specifically focusing on genes that disrupt osteoblast function, leading to monogenic bone disorders ^30–34^ (**Supplemental File 8**). We used a fisher’s exact test to measure the statistical significance of the representation of genes with associated mouse knockout bone phenotypes and monogenic bone disease in each module. We then applied a Bonferroni correction to correct for testing the enrichment of all 13 or 26 modules and applied a significance cutoff of 0.05 to the adjusted p-values.

### Clustering analysis in osteoblast differentiation gene expression data

We investigated the expression profiles of all purple module genes in the context of differentiation. Using gene expression data from osteoblasts throughout differentiation (Series GSE54461), we used k-means clustering to identify differentiation-related transcriptional programs in the purple module. We tested k = 1:5, and found two robust clusters of genes within the purple module. Enrichment analysis of the two clusters in all function categories were conducted as described above.

### Epigenetic enrichment analysis for LDS BMD GWAS associations

For BMD GWAS lead SNP (and proxies with LD >= 0.7) overlapping an LDS gene (n = 84), GenomicRanges ^66^ was used to calculate the proportion of lead SNPs overlapping regions marked by epigenetic modifications, including H3K4me1, H3K4me2, H3K4me3, H3K9me3, H3K27ac, H3K27me3, H3K26me3, H3K79me2 and H4K20me1, and histone H2AZ from the Roadmap Project 45. Using the GenomicRanges function findOverlaps(), we quantified the overlap between the LDS-associated lead SNPs and each epigenetic mark. To assess the enrichment of this overlap, we compared against 1000 sets of control SNPs (n = 84). We chose sets of control SNPs that were within +/-20% of the mean distance from a transcription start site for the BMD GWAS lead SNPs, and within +/-20% of the mean minor allele frequency of the BMD GWAS lead SNPs. P-values were calculated by taking the proportion of random sets of SNPs with a more extreme enrichment in the tail of the distribution with which we are comparing our experimental proportion. If the experimental proportion is more extreme than any measured random set, the p-value is reported as < 1×10_−3_. This same procedure was used to evaluate the tissue specificity for each mark. For each mark, the overlap with the LDS BMD SNP set and the 1000 random SNP sets were computed and the ratio between the proportion of overlapping LDS BMD SNPs and the mean proportion of overlapping random SNPs was computed. Higher ratios indicated greater enrichment of the LDS BMD SNPs over random SNPs with a given mark in a given tissue.

### Colocalization analysis

For each gene in the LDS that overlapped a BMD GWAS association from the Morris *et al.* study, eQTL from all GTEx tissues were identified ^10, 46^. Using the coloc package, we assessed the potential for colocalization between the QTL for BMD and the proximal cis-eQTL ^68^. Two associations were considered to colocalize if the posterior probability of hypothesis four (PPH4), which is the probability of colocalization, is > 0.7. The RACER package to plot the two associations in a mirrorPlot ^69^.

### Mouse phenotype statistical comparisons

Using the International Mouse Phenotyping Consortium (IMPC) database, we identified genes from the LDS that had eQTL that colocalized with BMD QTL and exhibited a difference in BMD when knocked out in mouse ^27^. Using the PhenStat package, we analyzed the differences between control and knockout animals using a mixed model framework ^70^. The specific equation used for each analysis are in **Supplemental File 15**.

### Network Topology Analysis

A t-test was used to compare the module membership of the four causal genes and the remainder of the LDS genes and the connectivity of the LDS genes overlapping a BMD GWAS locus as opposed to those that do not. A linear model was used to assess the relationship between gene connectivity and gene correlation with *in vitro* mineralization.

## Acknowledgements

Research reported in this publication was supported by the National Institute of Arthritis and Musculoskeletal and Skin Diseases of the National Institutes of Health under Award Number R01AR064790 to C.L.A-B. and C.R.F. O.L.S. was supported by a Wagner Fellowship from the University of Virginia. The Genotype-Tissue Expression (GTEx) Project was supported by the Common Fund of the Office of the Director of the National Institutes of Health, and by NCI, NHGRI, NHLBI, NIDA, NIMH, and NINDS. The data used for the analyses described in this manuscript (v7) were obtained from: the GTEx Portal on 01/11/2018. The International Mouse Phenotyping Consortium is partially funded by the NIH Knockout Mouse Programme (KOMP) project and the IMPC informatics and the data portal are supported by NIH grant U54 HG006370.

