## Supplemental Figures for "Identification of a core module for bone mineral density through the integration of a co-expression network and GWAS data"

**Supplemental Figure 1**. *In vitro* mineralization varied across the 42 strains of CC mice. Osteoblast cultures were stained ten days post-induction of differentiation with alizarin red stain. Inset images are examples of high mineralizing (strain = IL16768) and low mineralizing (strain = OR5306) strains.

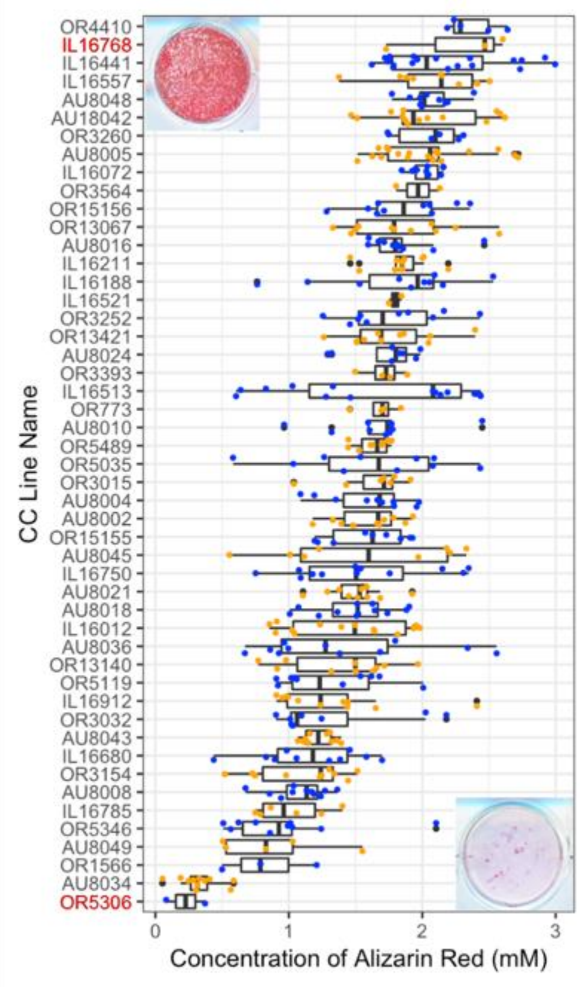

**Supplemental Figure 2**. The purple module eigengene was correlated with *in vitro* mineralization across the population.

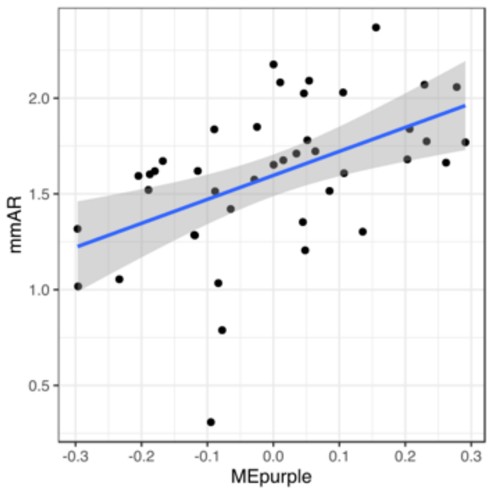

**Supplemental Figure 3**. More highly interconnected purple module genes (A) are more likely to overlap GWAS associations and (B) have patterns of expression more highly correlated with in vitro mineralization.

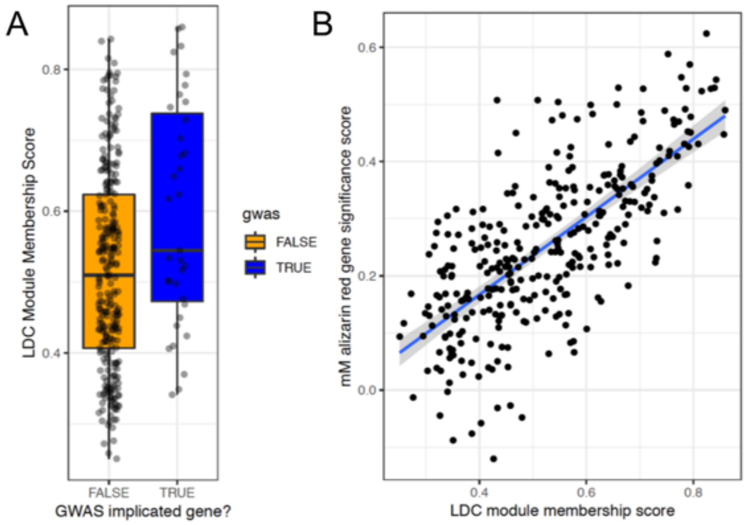
